# ST-DISTAL: Dual-Branch Graph Convolution with Distributional Alignment for Cell-Type Deconvolution

**DOI:** 10.64898/2026.09.27.754868

**Authors:** Abrar Rahman Abir, Muhtasim Noor Alif, Wei Zhang

## Abstract

Spatial transcriptomics enables high-throughput gene expression profiling while preserving spatial information, offering valuable insights into tissue architecture and cellular organization. Methods that achieve single-cell or subcellular spatial resolution typically rely on predefined gene panels, which limits genome-wide discovery. In contrast, sequencing-based spatial transcriptomics platforms provide broad transcriptome coverage but measure expression at lower spatial resolution, capturing mixtures of multiple cell types within each spatial location and thereby requiring accurate cell-type deconvolution. Many existing deconvolution methods do not jointly model molecular expression similarity and spatial context, fail to align predicted cell-type compositions with biological priors, and may produce spatially inconsistent results that do not reflect the smooth and structured organization of real tissues. We present ST-DISTAL, a dual-branch graph convolutional framework for cell-type deconvolution in spatial transcriptomics data. ST-DISTAL integrates conventional spatial graph convolution with spectral Chebyshev filtering through an adaptive attention mechanism, enabling the model to capture both local spatial dependencies and global structural patterns. The framework is trained using a composite objective that jointly enforces accurate cell-type proportion estimation, global distributional alignment with reference profiles, and spatial smoothness. Evaluations on three simulated benchmark datasets demonstrate that ST-DISTAL consistently outperforms state-of-the-art methods in both prediction accuracy and distributional alignment. Further validation on a real human embryonic heart dataset and a colorectal cancer dataset shows biologically plausible and spatially coherent cell-type organization.

## 1 Introduction

Tissues are mosaics of diverse cell types and states, and their spatial organization shapes how cells interact and function in both health and disease. Spatial transcriptomics technologies make it possible to measure gene expression while preserving this spatial context, producing maps of RNA abundance across a grid of spots laid over a tissue section (Moses and Pachter 2022; Ståhl et al. 2016; Vickovic et al. 2019). In most sequencing-based spatial transcriptomics methods, each spot captures RNA from a handful to dozens of cells, often mixing multiple cell types and states, and therefore yields only a bulk-like average (Rodriques et al. 2019; Eng et al. 2019; Lubeck et al. 2014; Moffitt et al. 2018). As a result, many spatial transcriptomics datasets contain high-dimensional but cell-type–mixed spots, which obscure both the true cellular composition and the precise location of specific subtypes (Cable et al. 2022; Ma and Zhou 2022). This leads to a central challenge: deconvolution, the task of determining which cell types are present at each spatial location and in what fractions, using only the mixed expression profiles observed in the spots (Li and Luo 2024; Chen et al. 2025b). Solving this problem is important because it allows spatial variation in gene expression to be understood in terms of underlying cell types and states rather than opaque mixtures. This, in turn, provides a solid foundation for downstream analyses such as cell-type–specific differential expression or spatially resolved pathway activity. In this way, accurate deconvolution transforms raw spatial transcriptomics measurements into a quantitative, cell-type–resolved view of the tissue, enabling systematic exploration and helping generate and test biological hypotheses.

Although emerging imaging-based spatial transcriptomics platforms can resolve gene expression at (sub)cellular resolution, they typically rely on targeted gene panels with limited coverage, which restricts their ability to characterize transcriptome-wide cellular states and compositions comprehensively (Wang et al. 2023). Because of this limitation, many computational methods use single-cell RNA-seq (scRNA-seq) data as a reference for spatial transcriptomics deconvolution (Li and Luo 2024; Yin et al. 2024; Zhou et al. 2023). scRNA-seq provides high-resolution, cell-level expression profiles with broad transcriptome coverage, capturing the transcriptional signatures of diverse cell types and states (Stuart and Satija 2019; Heumos et al. 2023). By using scRNA-seq–derived profiles as a reference, deconvolution methods can better constrain the inherently ill-posed inverse problem of unmixing spot-level expression, improving the identifiability of cell-type proportions and reducing reliance on unsupervised assumptions about the underlying cellular composition. Existing approaches span a wide range, from regression-based and probabilistic statistical models that infer cell-type proportions through likelihood-based estimation, to modern deep learning frameworks that directly predict compositional vectors using neural networks. However, benchmarking studies report substantial variability in performance across datasets and methods (Cable et al. 2022; Andersson et al. 2020).

Several challenges continue to limit progress in spatial deconvolution. At the data level, spatial transcriptomics measurements contain high technical noise and dropout, leaving many transcripts undetected and producing sparse expression profiles that obscure true cell-type signals (Rao et al. 2021; Shang and Zhou 2022). Methods that treat each spot independently and ignore spatial context can exacerbate this problem, because spatial continuity across neighboring spots carries valuable information that can help resolve mixed signals and improve biological plausibility (Li and Luo 2024; Long et al. 2023). On the modeling side, deep learning approaches, although increasingly adopted, often underperform in benchmarks and struggle to generalize across datasets (Yin et al. 2024). A major limitation is that their loss functions rarely enforce alignment between predicted cell-type proportions and distributions derived from single-cell references, which can lead to biased or inconsistent estimates. In addition, spatially adjacent spots tend to share similar gene expression patterns and cell-type compositions (Tu et al. 2025; Xia et al. 2019), but without explicit spatial regularization, models may produce abrupt and biologically implausible transitions across space. Addressing these interconnected issues is essential for accurate, robust, and scalable spatial transcriptomics analysis. To this end, we propose a new deep learning framework, **ST-DISTAL**, that performs accurate and spatially coherent cell-type deconvolution for spatial transcriptomics. Our main contributions are as follows:

1. We introduce a novel dual-branch graph convolutional layer that synergistically integrates standard spatial graph convolution with spectral Chebyshev polynomial convolution, adaptively fused through a learnable attention mechanism.
2. We employ a principled semi-supervised learning strategy combining distribution alignment via optimal transport with graph Laplacian regularization for spatial smoothness. These objectives encourage biologically plausible cell-type compositions that respect both global distributional structure and local tissue organization.
3. We achieve state-of-the-art performance on three simulated spatial transcriptomics benchmark datasets through comprehensive comparisons with recent methods, and further demonstrate the robustness and biological plausibility of ST-DISTAL via noise perturbation analyses, spatial domain recovery, and evaluation on a real human embryonic heart dataset.

## 2 Results

We compare ST-DISTAL with 13 recent state-of-the-art deconvolution methods across three spatial transcriptomics datasets, evaluating distributional alignment using Jensen–Shannon divergence (JSD), prediction error using root mean squared error (RMSE), and cell-type ranking accuracy using Spearman’s correlation. Detailed descriptions of the evaluation metrics are provided in the Supplementary Material. We further assess spatial domain recovery and robustness to noise through controlled Gaussian perturbation experiments, conduct ablation studies to quantify the contribution of individual model components, and validate biological plausibility using a real human embryonic heart spatial transcriptomics dataset.

### 2.1 Datasets

We evaluated ST-DISTAL on three publicly available single-cell spatial transcriptomics datasets from different platforms: seqFISH+ (Eng et al. 2019), seqFISH (Lubeck et al. 2014), and MER-FISH (Moffitt et al. 2018), selected for their high-resolution spatial coordinates and ground-truth cell-type annotations. Following Li and Luo (2024), we simulated spot-based measurements by aggregating single-cell expression profiles within square pixel grids, yielding paired spot-level data for deconvolution and single-cell profiles for evaluation.

The seqFISH+ dataset (mouse somatosensory cortex) profiles 10,000 genes across 315 cells and 4 cell-types, from which we generated 59 spots using 200 *×* 200 pixel grids. The seqFISH dataset (mouse embryo) contains 16,974 cells, 351 genes, and 4 cell-types, yielding 179 spots using 0.5 *×* 0.5 regions. The MERFISH dataset (mouse preoptic region) comprises 896,835 cells, 161 genes, and 7 cell-types, resulting in 23,097 spots using 50 *×* 50 pixel grids. Rare cell-types (< 3%) and low-coverage spots were excluded.

In addition, we evaluated ST-DISTAL on a human embryonic heart spatial transcriptomics dataset (Asp et al. 2019), generated using the original Spatial Transcriptomics technology, containing 210 spatial spots, 15 cell-types, and 13,467 genes. As spot-level ground truth is unavailable, this dataset is used to assess biological plausibility and spatial coherence.

For training, pseudo-spots were split into training and validation sets for supervision and early stopping. Real spots were included as unlabeled nodes in the graph and updated jointly with pseudo-spots via message passing, without introducing additional supervision.

### 2.2 Baselines

To ensure rigorous and comprehensive benchmarking, we selected 13 of the most recent state-of-the-art baselines: STdGCN (Li and Luo 2024), SpatialPrompt (Swain et al. 2024), NLSDeconv (Chen et al. 2025b), SDePER (Liu et al. 2024), CLPLS (Mo et al. 2025), QR-SIDE (Dong et al. 2025), NODE (Wang et al. 2025), LETSmix (Zhan et al. 2025), SpatialcoGCN (Yin et al. 2024), RCTD (Cable et al. 2022), Tangram (Biancalani et al. 2021), Cell2location (Kleshchevnikov et al. 2022) and SpatialCTD (Ding et al. 2024). All baseline implementations and benchmarking protocols were taken from the authors’ original code repositories or official publications whenever available. To ensure fairness, all models were run on the same hardware with identical data splits and we used the default hyperparameters exactly as provided by the authors. All experiments are conducted on a cluster equipped with four NVIDIA RTX A4500 GPUs. Hyperparameters for ST-DISTAL were optimized using Ray Tune (Liaw et al. 2018) with an ASHA scheduler (Li et al. 2020). We report the final hyperparameters in the Supplementary Material.

### 2.3 ST-DISTAL Achieves State-of-the-Art Prediction Accuracy

We assess cell-type deconvolution performance using RMSE, a standard metric that captures the average difference between predicted and true cell-type proportions for each spatial spot. Lower RMSE reflects higher accuracy and closer alignment with ground-truth composition. Table 1 presents the RMSE of ST-DISTAL and baselines across the three benchmark datasets. On seqFISH+, ST-DISTAL achieves the lowest RMSE (0.0689), outperforming the next best method, SpatialcoGCN (0.0712), by 3.23%. The improvement margin over the third best, CLPLS (0.0721), is 4.44%. For the seqFISH dataset, ST-DISTAL attains the best RMSE (0.1674), with improvements of 5.9% and 17.5% over SpatialCTD (0.1779) and STdGCN (0.2029), respectively. These results highlight the robustness and predictive superiority of ST-DISTAL on diverse single-cell resolution data. For the MERFISH dataset, ST-DISTAL (0.1293) obtains the best performance, outperforming all the baselines by a substantial margin (e.g., 5.96% over STdGCN and 7.58% over SpatialcoGCN). Notably, many existing methods exhibit markedly higher RMSE, even by more than an order of magnitude, demonstrating the superiority of ST-DISTAL. All experiments were repeated ten times. Statistical significance was assessed using the Wilcoxon rank sum test, with ST-DISTAL achieving significantly better performance than all baseline methods (*p*-value *<* 0.05). By explicitly enforcing both distributional alignment via optimal transport and spatial coherence through Laplacian regularization, ST-DISTAL leverages biologically grounded objectives that lead to superior deconvolution accuracy. This principled approach enables the model to capture the intrinsic diversity and spatial organization of cell-types, resulting in state-of-the-art performance across benchmarks.

**Table 1:** Comparison of RMSE (↓) and JSD (↓) scores across all methods and datasets. The lowest RMSE and JSD for each dataset is bolded.

| Method | RMSE |  |  | JSD |  |  |
| --- | --- | --- | --- | --- | --- | --- |
|  | seqFISH+ | seqFISH | MERFISH | seqFISH+ | seqFISH | MERFISH |
| ST-DISTAL | **0.0689** | **0.1674** | **0.1293** | **0.0134** | **0.0729** | **0.0180** |
| STdGCN | 0.0736 | 0.2029 | 0.1375 | 0.0139 | 0.0765 | 0.0798 |
| SpatialPrompt | 0.4514 | 0.3264 | 0.1562 | 0.3778 | 0.2275 | 0.0724 |
| NLSDeconv | 0.0744 | 0.2131 | 0.1504 | 0.0148 | 0.0768 | 0.0864 |
| SDePER | 0.0779 | 0.2090 | 0.1544 | 0.0150 | 0.0826 | 0.0929 |
| CLPLS | 0.0721 | 0.2074 | 0.1532 | 0.0145 | 0.0805 | 0.0808 |
| QR-SIDE | 0.0746 | 0.2095 | 0.1538 | 0.0153 | 0.0819 | 0.0906 |
| NODE | 2.0670 | 4.2076 | 3.9810 | 1.4381 | 3.3469 | 5.6195 |
| LETSmix | 0.4536 | 0.3497 | 0.1741 | 0.3729 | 0.2484 | 0.0897 |
| SpatialcoGCN | 0.0712 | 0.2104 | 0.1399 | 0.0154 | 0.0854 | 0.0828 |
| RCTD | 0.1082 | 0.2579 | 0.2118 | 0.0293 | 0.1213 | 0.0396 |
| Cell2location | 0.1385 | 0.3012 | 0.2784 | 0.0352 | 0.1594 | 0.0488 |
| Tangram | 0.1914 | 0.3688 | 0.3236 | 0.0578 | 0.2147 | 0.0693 |
| SpatialCTD | 0.1609 | 0.1779 | 0.1447 | 0.0626 | 0.0840 | 0.0294 |

### 2.4 ST-DISTAL Excels in Accurate Cell-Type Distribution Alignment

We report the Jensen–Shannon Divergence (JSD) across all methods and datasets to evaluate the alignment between predicted and ground-truth cell-type distributions. JSD quantifies the similarity between two distributions, with lower values indicating a closer match between predicted and true cell-type proportions within each spatial spot. As shown in Table 1, ST-DISTAL consistently achieves the lowest JSD values on all three benchmark datasets, indicating highly precise alignment of predicted cell-type distributions. On seqFISH+, ST-DISTAL attains a JSD of 0.0134, outperforming STdGCN (0.0139) and CLPLS (0.0145) by 3.6% and 7.6%, respectively. For seqFISH, ST-DISTAL yields a JSD of 0.0729, offering improvements of 4.7% and 5.1% over the next-best methods, STdGCN (0.0765) and NLSDeconv (0.0768). On MERFISH, ST-DISTAL achieves a JSD of 0.0180, surpassing SpatialCTD (0.0294) and RCTD (0.0396) by 38.8% and 54.5%, respectively. Collectively, these results confirm that ST-DISTAL provides the most accurate global distributional alignment, substantially narrowing the gap between predicted and true cell-type compositions compared to all recent baselines.

### 2.5 ST-DISTAL Demonstrates Superior Cell-Type Ranking Concordance

We evaluated cell-type ranking concordance performance using Spearman’s rank correlation coefficient as shown in Figure 1. This metric quantifies how well the predicted rankings of cell-type proportions match the true ground-truth rankings, capturing both the order and monotonicity of predictions. Higher values indicate more faithful reconstruction of relative cell-type abundances. Across all datasets, ST-DISTAL consistently achieves top or near-top Spearman’s rank correlation, highlighting its ability to preserve the underlying cell-type order. On the seqFISH+ dataset, ST-DISTAL attains the highest Spearman’s correlation of 0.8324, outperforming the second-best method SDePER (0.8009) by 3.9%, and STdGCN (0.7853) by 6.0%. For the seqFISH dataset, ST-DISTAL again leads with a score of 0.8079, improving upon STdGCN (0.7854) by 2.9% and SDePER (0.7747) by 4.3%. On the MERFISH dataset, ST-DISTAL achieves the best performance (0.7128), with SpatialCTD (0.6810) as the closest competitor. All experiments were repeated ten times, with ST-DISTAL significantly outperforming all baselines (*p*-value *<* 0.05, Wilcoxon rank sum test).

**Figure 1.**
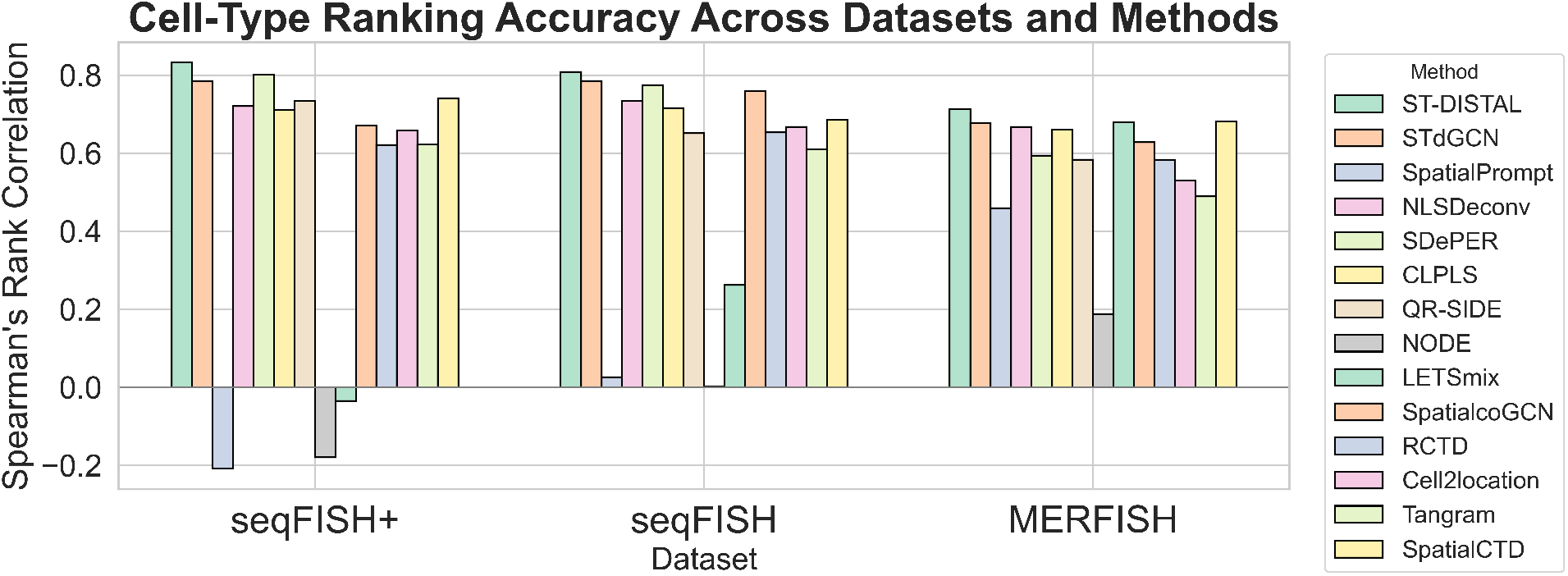
Comparison of Spearman’s rank correlation coefficients for cell-type composition prediction across three datasets. ST-DISTAL achieves the highest concordance in seqFISH+ and seqFISH, and remains competitive in MERFISH.

### 2.6 ST-DISTAL Yields High-Fidelity Spatial Domain Recovery

To assess ST-DISTAL’s ability to preserve biologically meaningful spatial domains, we performed a quantitative domain recovery analysis on the seqFISH dataset. Each spot was labeled by its ground-truth cell-type composition, while predicted cell-type proportions from all methods were clustered via Agglomerative Clustering to infer domain assignments for direct comparison with the reference. We evaluated performance using the Adjusted Rand Index (ARI) and Normalized Mutual Information (NMI), two widely adopted metrics for quantifying the concordance between predicted and ground-truth clustering.

As presented in Figure 2, ST-DISTAL achieves the highest ARI (0.6094) and NMI (0.6562) among all compared methods, outperforming strong baselines such as STdGCN (ARI: 0.5521, NMI: 0.6120) and SpatialcoGCN (ARI: 0.5547, NMI: 0.6153). Figure 3 visualizes cell-type compositions for each spot across all methods using pie charts, highlighting ST-DISTAL’s strong agreement with the ground truth. These results confirm that ST-DISTAL provides accurate cell-type deconvolution while effectively preserving spatial domain architecture, essential for reliable downstream analysis.

**Figure 2.**
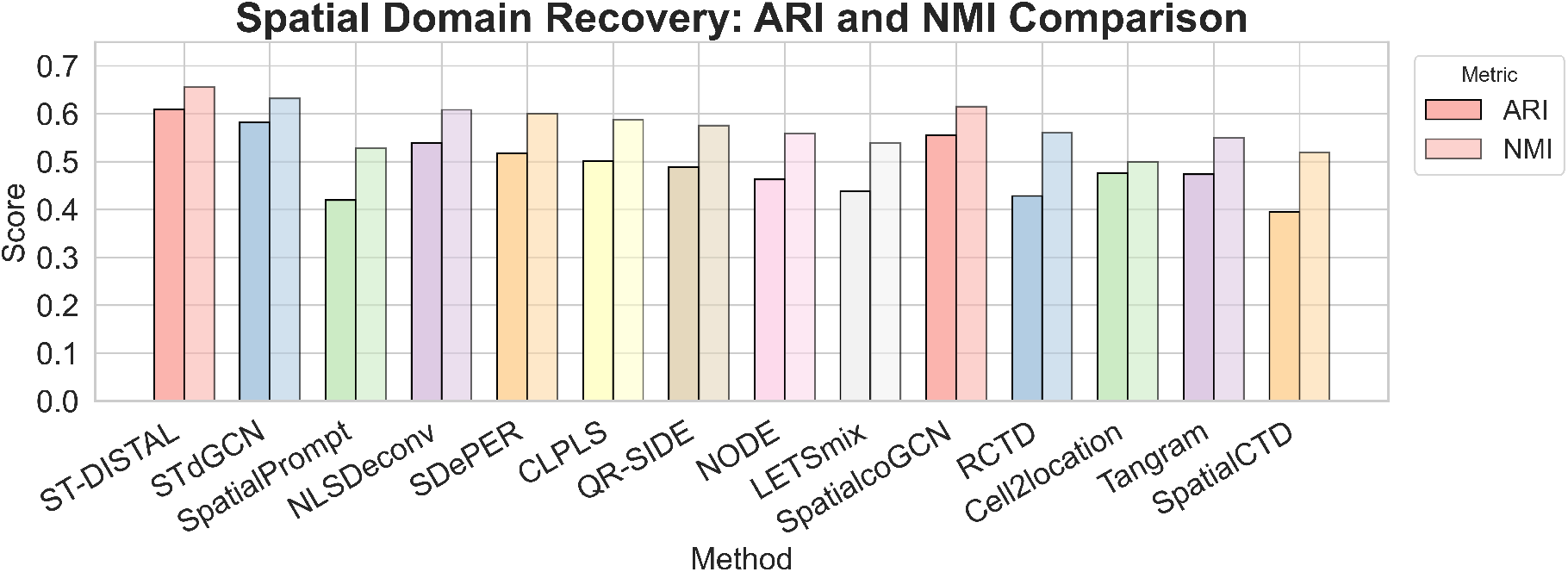
Quantitative comparison of spatial domain recovery by all methods, measured by Adjusted Rand Index (ARI) and Normalized Mutual Information (NMI). ST-DISTAL achieves the highest ARI and NMI, indicating superior fidelity in preserving the true spatial domain structure.

**Figure 3.**
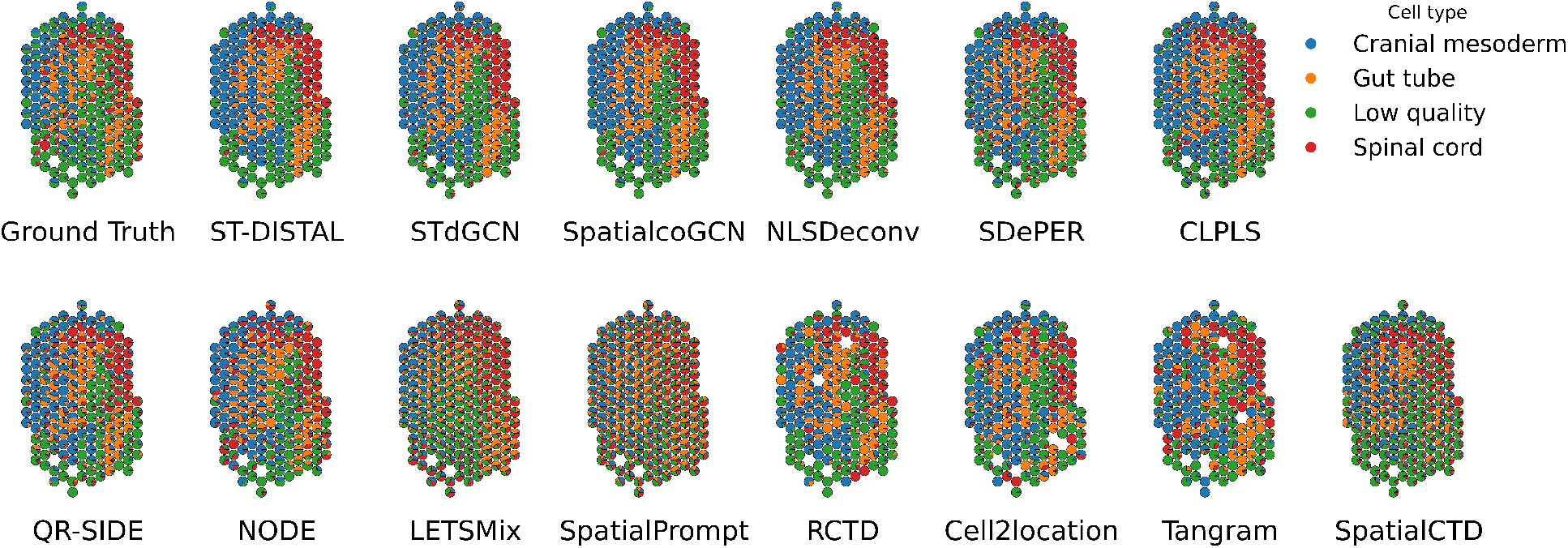
Comparison of spatial cell-type composition across all methods. Each subplot visualizes the cell-type proportions at every spatial spot as pie charts for the seqFISH dataset. ST-DISTAL most accurately reconstructs the ground-truth spatial organization, while other baselines display varying levels of spatial and compositional distortion.

### 2.7 ST-DISTAL is Resilient to Input Feature Noise Perturbation

To assess the stability of ST-DISTAL under noise perturbation, we performed a controlled robustness analysis by injecting Gaussian noise into the latent embeddings for both real and pseudo-spots prior to dual-branch GCN processing. This simulates common sources of technical or biological variability that may affect the reliability of spatial transcriptomics data in practical scenarios. Figure 4 presents the trend of key evaluation metrics on the seqFISH+ dataset, measured as the standard deviation of the injected noise increases. We observe that the predictive accuracy of ST-DISTAL remains consistently high for moderate levels of feature perturbation, with only a gradual degradation in performance as noise becomes severe. Notably, the model maintains competitive cell-type deconvolution performance even when subjected to substantial input noise, highlighting its reliability for practical applications where measurement errors are unavoidable.

**Figure 4.**
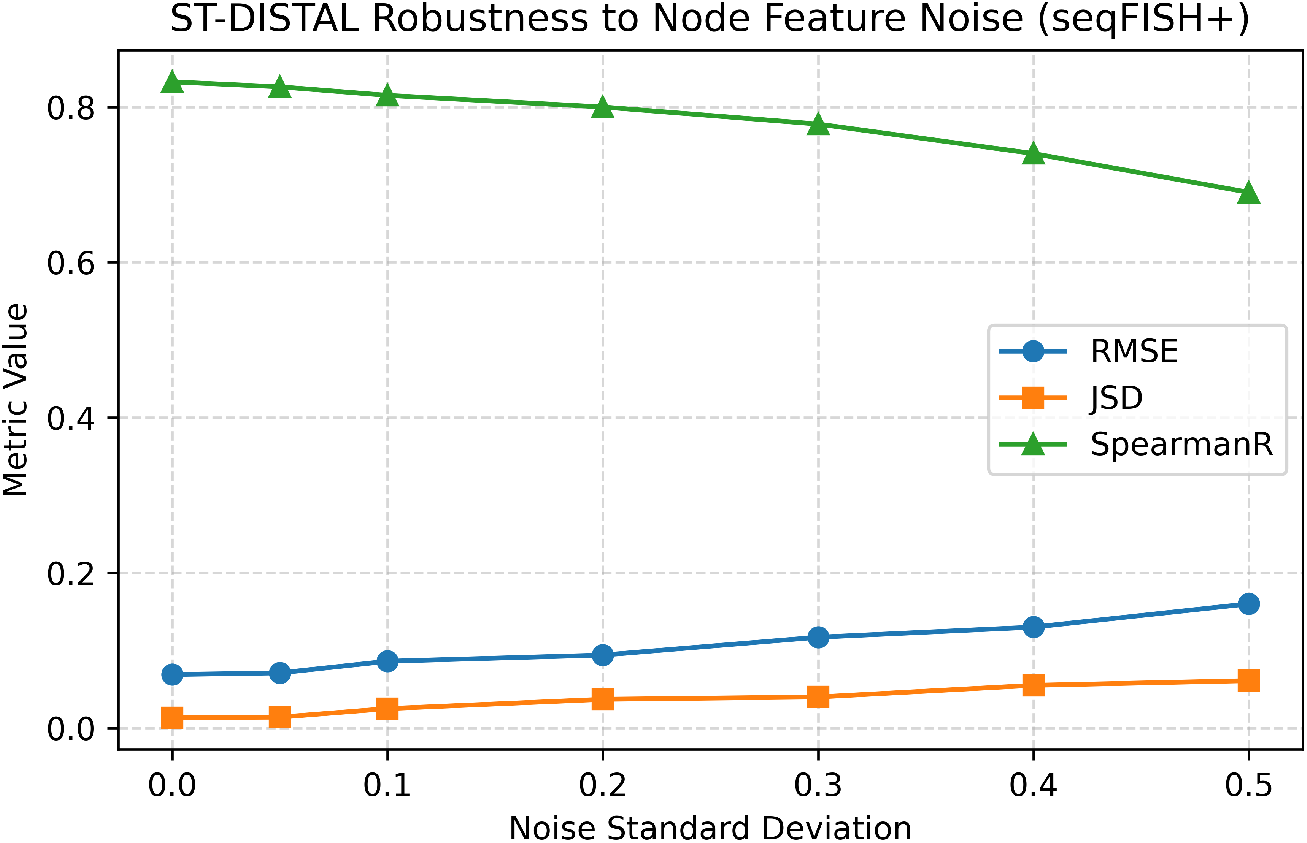
Robustness of ST-DISTAL to input noise. Performance (JSD, RMSE, Spearman’s correlation) on the seqFISH+ dataset with increasing Gaussian noise in node features. ST-DISTAL remains stable across noise levels, showing strong resilience to feature perturbations.

### 2.8 ST-DISTAL Recovers Biologically Meaningful Cell-Type Composition from Real Embryonic Heart Spatial Transcriptomics Data

We analyzed a human embryonic heart spatial transcriptomics dataset collected at approximately 6.5 post-conceptional weeks (Asp et al. 2019). The data were generated using the original Spatial Transcriptomics technology, where gene expression is measured at spot resolution, with each spot capturing transcripts from multiple cells rather than individual cells. As a result, direct identification of cell-type composition from the spatial data alone is not possible. Single-cell transcriptomic analysis of the developing human heart has shown that ventricular cardiomyocytes expand rapidly during early developmental stages and constitute the dominant cardiomyocyte population. In contrast, atrial cardiomyocytes exhibit greater diversification and expansion at later stages of heart development (Asp et al. 2019; Cui et al. 2019). Motivated by this biological observation, we applied ST-DISTAL to infer spot-level cell-type compositions from the spatial transcriptomics data. Using reference cell-type gene signatures derived from scRNA-seq data, ST-DISTAL estimates the proportion of each cardiac cell type at every spatial location. The inferred cell-type composition, as shown in Figure 5, identified ventricular cardiomyocytes as the dominant population across the tissue section, yielding a biologically plausible representation of embryonic heart cellular organization. This experiment shows that ST-DISTAL recovers known biological properties of embryonic heart tissue from real spatial transcriptomics data.

**Figure 5.**
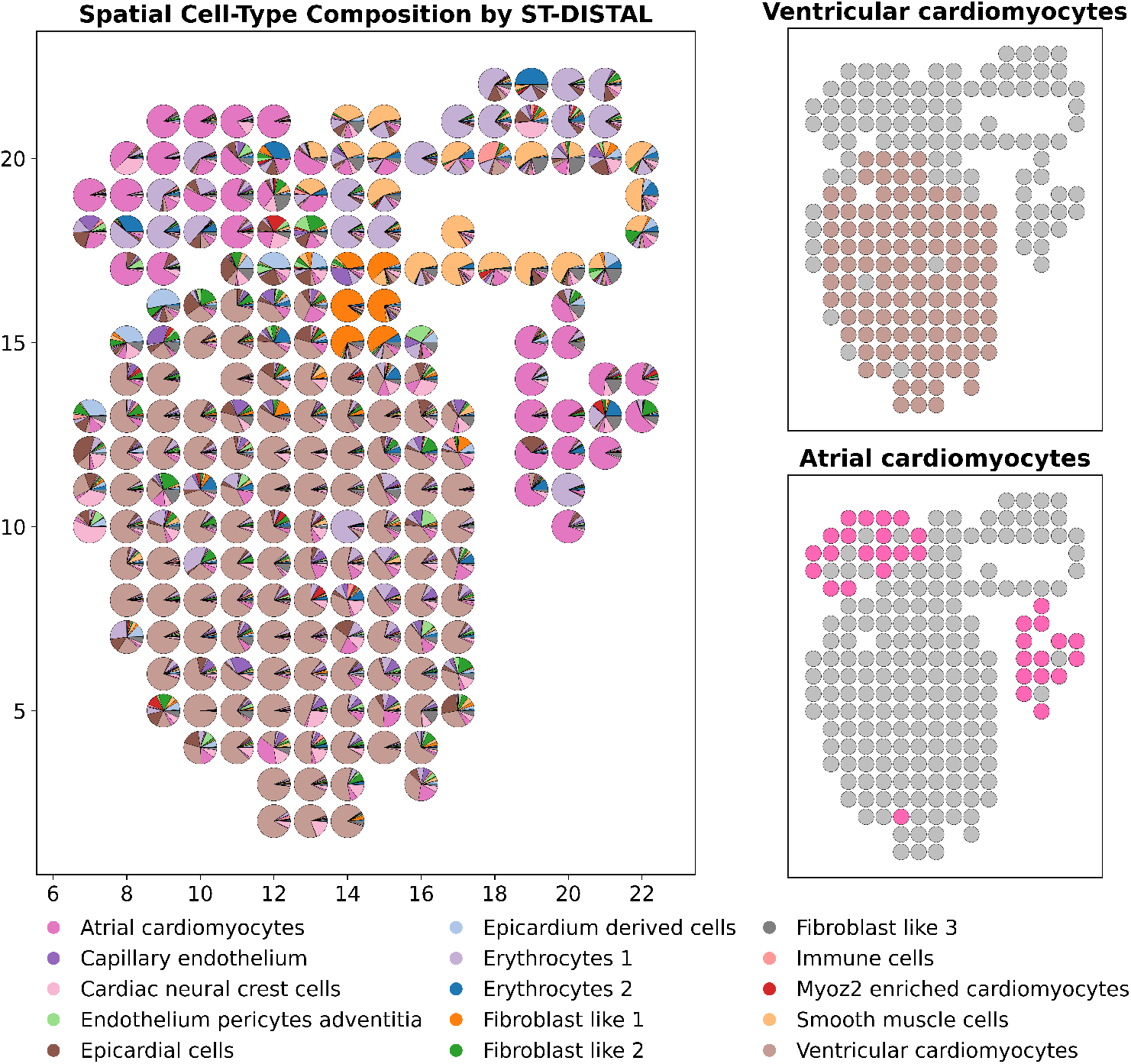
Spatial visualization of inferred cell-type composition in embryonic heart tissue. Each spot represents a spatial location and is shown as a pie chart of estimated cell-type proportions. Ventricular cardiomyocytes are inferred as the dominant population across the tissue, while atrial cardiomyocytes exhibit more spatially localized enrichment.

### 2.9 ST-DISTAL recovers tumor-proximal cellular organization in human col-orectal cancer

We evaluated whether ST-DISTAL recovers biologically plausible spatial organization in human colorectal cancer by analyzing its inferred cell-type proportions relative to computationally identified tumor regions (Fig. 6). Tumor regions were defined independently of ST-DISTAL from CRC-associated expression patterns, and the tissue surrounding each tumor boundary was divided into distance intervals, with the first 50 *µ*m treated as the primary tumor-proximal compartment following previous Visium HD CRC analyses (Oliveira et al. 2025). We then tested whether specific cell populations were preferentially enriched near the tumor and whether their abundance changed with increasing distance. Myeloid/macrophage cells showed the strongest enrichment within 50 *µ*m of the tumor boundary (ER = 1.489; 95% CI, 1.310–1.661), followed by T/NK cells (ER = 1.255; 95% CI, 1.123–1.397) (Fig. 6B,C). Myeloid/macrophage abundance progressively decreased from 0.183 at 0–50 *µ*m to 0.159 at 50–100 *µ*m, 0.149 at 100–200 *µ*m, and 0.121 beyond 200 *µ*m (Fig. 6D). Myeloid/macrophage cells were also the most abundant inferred immune population in the immediate tumor neighborhood, exceeding T/NK and B-cell proportions. These observations are consistent with previous high-definition spatial profiling of human CRC, which reported prominent macrophage accumulation and structured immune organization near tumor interfaces (Oliveira et al. 2025).

**Figure 6.**
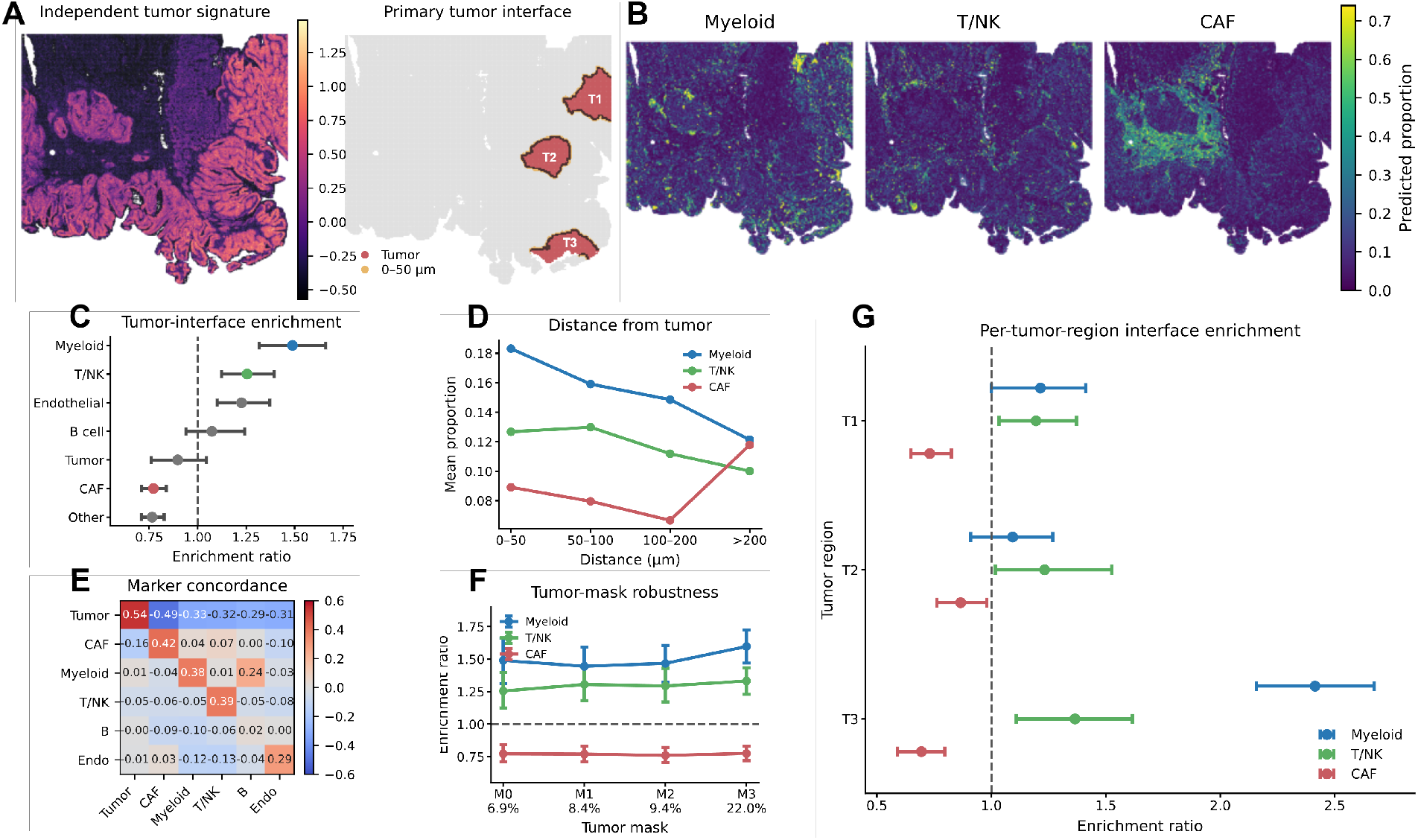
ST-DISTAL recovers tumor-proximal cellular organization in human colorectal cancer. **(A)** Independent tumor-signature map and primary tumor interface with three spatially distinct tumor regions (T1–T3). **(B)** Spatial distributions of inferred myeloid/macrophage, T/NK, and fibroblast/CAF proportions. **(C)** Cell-type enrichment within 0–50 *µ*m of the tumor boundary, with 95% spatial-bootstrap confidence intervals. **(D)** Mean cell-type proportions across increasing tumor-distance intervals. **(E)** Correlation between inferred cell-type proportions and canonical marker programs. **(F)** Tumor-mask sensitivity across M0–M3. **(G)** Per-region enrichment across T1–T3, showing reproducible myeloid/macrophage and T/NK enrichment. Dashed lines indicate no enrichment (ER = 1).

Fibroblast/CAF abundance exhibited a different, non-monotonic spatial pattern. Although CAFs were not globally enriched within the 0–50 *µ*m compartment relative to all more distant tissue (ER = 0.772), their mean abundance was higher immediately adjacent to the tumor than at 50–100 *µ*m and 100–200 *µ*m, before increasing again in distal fibroblast-rich regions (Fig. 6C,D). This pattern suggests that CAFs are not restricted to a narrow tumor boundary but instead occupy spatially heterogeneous stromal compartments across the tissue. Such heterogeneity is consistent with previous spatial studies of CRC showing distinct fibroblast/CAF-rich regions and variable localization of CAF-associated transcriptional programs (Chen et al. 2025a). Importantly, an independent analysis of the same H1-VM2JXXK Visium HD specimen demonstrated spatial correspondence between fibroblast-associated regions and established CAF markers, providing additional support for the heterogeneous stromal organization observed here (Chen et al. 2025a).

The inferred cell populations were further supported by concordance with their canonical transcriptional programs (Fig. 6E). Positive marker concordance was observed for tumor/epithelial (*ρ* = 0.541), fibroblast/CAF (*ρ* = 0.416), T/NK (*ρ* = 0.386), myeloid/macrophage (*ρ* = 0.384), and endothelial populations (*ρ* = 0.294). This agreement indicates that regions assigned higher proportions of a given cell type generally exhibited stronger expression of the corresponding marker program. Similar spatial agreement between cell-type assignments and canonical markers, including tumor- and fibroblast-associated genes, has been used as supporting biological validation in previous Visium HD CRC analyses (Oliveira et al. 2025; Chen et al. 2025a).

The macrophage-associated pattern was also robust to alternative tumor definitions. Across four increasingly permissive tumor masks (M0-M3), myeloid/macrophage enrichment remained between 1.445- and 1.597-fold, while T/NK enrichment remained consistently above 1 (Fig. 6F). Importantly, the same pattern was reproduced across the three spatially distinct tumor regions identified in the tissue (Fig. 6A,G). Myeloid/macrophage abundance was enriched around each tumor region, with the strongest enrichment surrounding T3, while T/NK cells were also consistently elevated. In contrast, CAF enrichment remained below one across the three interfaces, reinforcing the conclusion that fibroblast/CAF organization is spatially heterogeneous rather than uniformly concentrated at tumor boundaries. Together, these results indicate that ST-DISTAL recovers a reproducible macrophage-dominant tumor-proximal microenvironment, together with associated lymphoid enrichment and heterogeneous stromal organization, in agreement with established spatial features of human CRC (Oliveira et al. 2025; Chen et al. 2025a).

### 2.10 Ablation Study

To elucidate the contribution of major components in ST-DISTAL, we conducted a series of ablation experiments as shown in Table 2. Replacing the attention-based fusion with simple averaging of the GCN and spectral branches resulted in a modest performance decrease, confirming the importance of adaptive branch weighting. Excluding the spectral branch and using only the standard GCN further degraded performance, highlighting the benefit of dual-pathway feature extraction. In addition, ablation of either the OT loss or the Laplacian regularization term individually degraded performance, suggesting complementary benefits. Joint removal produced the largest drop, underscoring the necessity of distributional alignment and spatial smoothness. All ablation experiments were repeated ten times, and statistical significance was assessed using Wilcoxon rank sum test. Collectively, these findings demonstrate that each module in ST-DISTAL is essential for achieving optimal performance.

**Table 2:** Ablation Study of ST-DISTAL. Lower JSD/RMSE and higher SpearmanR are better. * indicates statistically significant difference (*p*-*value <* 0.05) from ST-DISTAL.

| Variant | seqFISH+ |  |  | seqFISH |  |  | MERFISH |  |  |
| --- | --- | --- | --- | --- | --- | --- | --- | --- | --- |
|  | JSD | RMSE | SpearmanR | JSD | RMSE | SpearmanR | JSD | RMSE | SpearmanR |
| ST-DISTAL | **0.0134** | **0.0689** | **0.8324** | **0.0729** | **0.1674** | **0.8079** | **0.0180** | **0.1293** | **0.7128** |
| w/o attention fusion | 0.0139 | 0.0720* | 0.8241 | 0.0743 | 0.2021 | 0.7957 | 0.0795 | 0.1340 | 0.6991 |
| w/o spectral branch | 0.0146* | 0.0763* | 0.8132* | 0.0758* | 0.2095* | 0.7823* | 0.0811* | 0.1397* | 0.6764* |
| w/o OT loss | 0.0147* | 0.0746* | 0.8189* | 0.0756* | 0.1987* | 0.7912* | 0.0801* | 0.1371* | 0.6849* |
| w/o Laplacian loss | 0.0142* | 0.0729* | 0.8235* | 0.0748* | 0.1913* | 0.7968* | 0.0786* | 0.1348* | 0.6917* |
| w/o OT+Laplacian loss | 0.0154* | 0.0784* | 0.8015* | 0.0772* | 0.2158* | 0.7691* | 0.0830* | 0.1442* | 0.6622* |

## 3 Discussion

In this work, we introduce ST-DISTAL, a dual-branch GNN for spatial transcriptomics cell-type deconvolution that combines adaptive dual-branch graph convolution with distributional alignment. Extensive experiments demonstrate that ST-DISTAL consistently outperforms recent baselines across multiple benchmarks, achieving lower prediction error, improved distributional alignment, and superior cell-type ranking concordance. Furthermore, ablation studies and noise sensitivity analysis confirm the importance of each model component and the resilience of our model to input noise. As a promising future direction, we plan to extend ST-DISTAL to leverage multi-omics spatial datasets for integrated cell-type and state inference.

## 4 Methods

### 4.1 Problem Formulation

Let 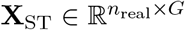 denote the spatial transcriptomics gene expression matrix consisting of *n*_real_ spatial spots and *G* genes. Each spot is assumed to contain a mixture of *C* cell-types. For each spot *i*, we aim to infer a cell-type proportion vector 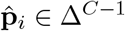, where *C* is the total number of cell-types and Δ^*C™*1^ is the (*C*™1)-dimensional probability simplex. The set of *C* cell-types is defined based on a reference scRNA-seq dataset with gene-matched, cell-type–annotated profiles. In simpler terms, given only the gene expression and spatial location of each spot, we aim to accurately estimate the proportion of each cell-type present at every spot in the tissue.

### 4.2 Overview of the ST-DISTAL Framework

ST-DISTAL performs cell-type deconvolution in spatial transcriptomics by leveraging both molecular similarity and spatial topology. Starting from a reference scRNA-seq dataset with annotated cell-types, we preprocess the data to remove technical variation and select informative marker genes. We then generate synthetic pseudo-spots by mixing randomly sampled single-cell profiles, providing supervised training examples with known cell-type proportions. Both real spatial spots and pseudo-spots are embedded into a shared low-dimensional space using these markers. In this space, we construct two graphs: an expression similarity graph linking spots by transcriptomic similarity, and a spatial proximity graph. Our dual-branch graph convolutional network (GCN) processes each graph independently, combining a standard graph convolution branch with a spectral Chebyshev polynomial branch, fused at each layer via attention. The resulting node embeddings from both graphs are concatenated and decoded by a multilayer perceptron to predict cell-type proportions. Finally, the model is optimized in a semi-supervised setting, where supervision is available only for pseudo-spots with known cell-type proportions. A Kullback–Leibler (KL) divergence loss is employed on these labeled pseudo-spots to directly compare predicted and reference cell-type distributions, making it a natural choice for proportion-valued outputs that lie on the probability simplex. However, accurate deconvolution also requires preserving biologically meaningful relationships across the tissue. Rather than relying solely on pointwise supervision, we introduce an optimal transport (OT) loss to align the global distribution of predicted proportions with that of the reference, explicitly accounting for similarities and transitions between cell types. This distribution-level constraint is particularly important in spatial transcriptomics, where cell-type compositions vary continuously across tissue rather than independently across spots. In parallel, a graph Laplacian regularization term encourages spatial smoothness by penalizing abrupt changes in predicted compositions between neighboring spots, reflecting the biological principle that neighboring tissue regions tend to share similar cellular organization. Together, these complementary losses enforce local accuracy, global distributional consistency, and spatial coherence.

Next, we briefly discuss the pseudo-spot simulation, graph construction, dual-branch graph convolutional network and the training objective. Table 3 contains the summary of the notations we use. Figure 7 shows the complete pipeline of ST-DISTAL.

**Table 3:** Summary of Notation.

| Symbol | Description |
| --- | --- |
| $n_{\text{real}}$ | Number of real spatial spots |
| $n_{\text{ps}}$ | Number of synthetic pseudo-spots |
| $n_{\text{sc}}$ | Number of reference single-cells |
| $N = n_{\text{real}} + n_{\text{ps}}$ | Total number of spots (real + pseudo-spots) |
| $G$ | Number of genes |
| $C$ | Number of cell-types |
| $d, d'$ | Dimensionality of low-dimensional embedding spaces |
| $\mathbf{X}_{\text{ST}} \in \mathbb{R}^{n_{\text{real}} \times G}$ | Spatial transcriptomics gene expression matrix |
| $\mathbf{X}_{\text{sc}} \in \mathbb{R}^{n_{\text{sc}} \times G}$ | Reference scRNA-seq expression matrix |
| $\tilde{\mathbf{X}}_{\text{real}} \in \mathbb{R}^{n_{\text{real}} \times d}$ | Embedded latent features of real spots |
| $\tilde{\mathbf{X}}_{\text{ps}} \in \mathbb{R}^{n_{\text{ps}} \times d}$ | Embedded latent features of pseudo-spots |
| $\mathbf{A}^{\text{expr}} \in \{0, 1\}^{N \times N}$ | Expression similarity graph adjacency matrix |
| $\mathbf{A}^{\text{spatial}} \in \mathbb{R}^{N \times N}$ | Spatial proximity graph adjacency matrix |
| $\mathbf{Z}^{\text{expr}} \in \mathbb{R}^{N \times d'}$ | Learned embeddings from expression graph |
| $\mathbf{Z}^{\text{spatial}} \in \mathbb{R}^{N \times d'}$ | Learned embeddings from spatial graph |
| $\mathbf{Z}^{\text{fused}} \in \mathbb{R}^{N \times 2d'}$ | Final fused node embedding |
| $\mathbf{P}^{\text{ref}} \in \mathbb{R}^{n_{\text{ps}} \times C}$ | Ground-truth pseudo-spot proportion matrix |
| $\hat{\mathbf{P}} \in \mathbb{R}^{N \times C}$ | Predicted cell-type proportion matrix for all spots |
| $\mathbf{p}_i \in \Delta^{C-1}$ | Ground-truth cell-type proportions for pseudo-spot $i$ |
| $\hat{\mathbf{p}}_i \in \Delta^{C-1}$ | Predicted cell-type proportions for spot $i$ |
| $\mathbf{M} \in \mathbb{R}^{N \times n_{\text{ps}}}$ | Optimal transport cost matrix |
| $\mathbf{T} \in \mathbb{R}^{N \times n_{\text{ps}}}$ | Optimal transport plan |

**Figure 7.**
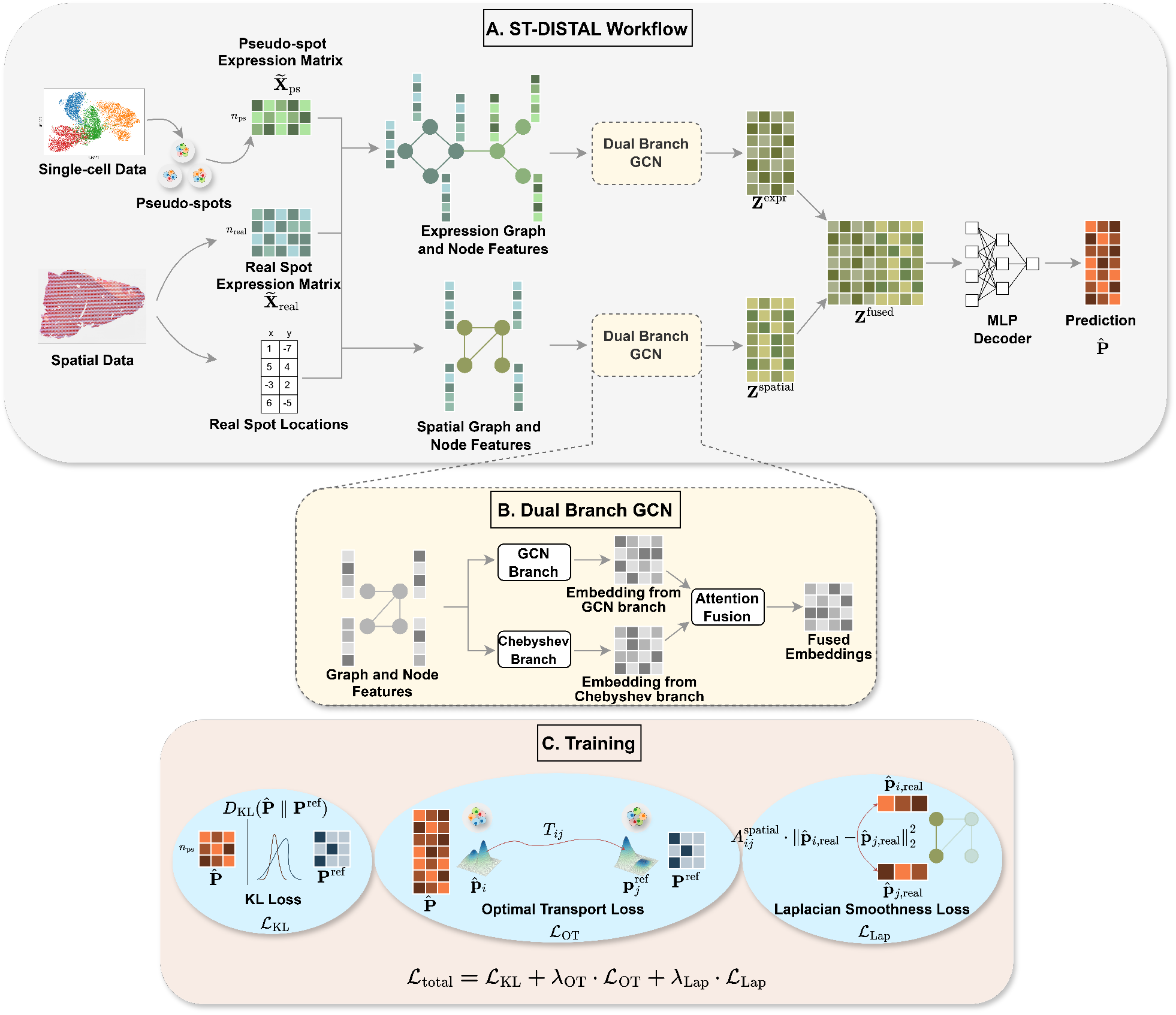
Overview of the ST-DISTAL framework. (A) The pipeline generates pseudo-spots and combines scRNA-seq reference and spatial transcriptomics data by constructing expression and spatial graphs, and encoding them via a dual-branch GCN to predict spot-level cell-type proportions. (B) Each GCN layer fuses standard spatial and Chebyshev spectral branches using an adaptive attention mechanism. (C) The model is trained with a composite loss: KL divergence on pseudo-spots, optimal transport for distribution alignment, and Laplacian regularization for spatial smoothness.

### 4.3 Pseudo-spot Simulation

Following (Li and Luo 2024), we simulate synthetic training samples, referred to as pseudo-spots, using a reference scRNA-seq dataset 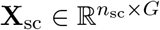, where each cell is annotated with a known label *y*_*j*_ ∈ *{*1, …, *C}*. These pseudo-spots act as supervised examples with known cell-type compositions, enabling model training in the absence of labeled spatial data.

Prior to simulation, we preprocess **X**_sc_ to reduce technical noise and highlight biological variation by selecting a set of marker genes and applying dimensionality reduction. Let *ϕ* : ℝ^*G*^ → ℝ^*d*^ denote the projection function into a shared *d*-dimensional embedding space. Both the spatial transcriptomics matrix 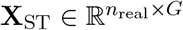 and pseudo-spot data are mapped via *ϕ* into the same latent space, yielding 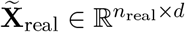 and 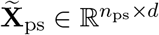.

To construct each pseudo-spot, we sample a small subset of cell types *C*_*i*_ ⊆ *{*1, …, *C}*, reflecting the limited diversity observed in real spatial spots. We then sample up to a predefined maximum number of single cells from the reference scRNA-seq dataset, restricted to cell types in *C*_*i*_, and aggregate their expression profiles to form a pseudo-spot. The number of cells contributed by each cell type is determined by the sampling process and may vary across pseudo-spots. A stratified sampling scheme is used to ensure that all cell types are adequately represented across the pseudo-spot training set. The resulting cell-type proportions **p**_*i*_ ∈ Δ^*C™*1^ are recorded as supervisory labels.

### 4.4 Graph Construction

We construct two complementary graphs over the combined set of real and pseudo-spots: an expression similarity graph and a spatial proximity graph. Using the previously defined embeddings 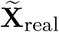 and 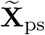, we define the full node set as *V* = *V*_real_ ∪ *V*_ps_, where each node corresponds to a spot in the spatial transcriptomics tissue or a synthetically generated pseudo-spot.

The expression graph *G*_expr_ = (*V, E*_expr_) encodes transcriptional similarity across all nodes in the joint embedding space. It comprises three subgraphs: a *real-to-real* subgraph between spatial real spots, a *pseudo-to-pseudo* subgraph among synthetic pseudo-spots, and a bipartite *real-to-pseudo* subgraph linking the two. For each subgraph, we compute pairwise cosine distances in the latent space and construct edges using the mutual nearest neighbor (MNN) criterion. Specifically, an edge is formed between two nodes *a* and *b* if *a* is among the top-*k* nearest neighbors of *b*, and *b* is also among the top-*k* nearest neighbors of *a*. The resulting graph *G*_expr_ enables expression-driven message passing between real and pseudo nodes, allowing supervision from labeled pseudo-spots to propagate into the spatial domain. The corresponding adjacency matrix *A*^expr^ is binary and symmetric.

The spatial graph *G*_spatial_ = (*V, E*_spatial_) models the geometric structure of the tissue. To construct this graph, we use the physical coordinates of each spot on the tissue section to compute pairwise distances. Edges are then created between spots that fall within a specified spatial radius, reflecting local physical proximity. The adjacency matrix *A*^spatial^ is weighted, with edge weights inversely related to the spatial distance, thereby promoting smoothness across nearby tissue regions. Conceptually, the spatial graph is defined using only real spots based on their coordinates. For joint processing, the spatial graph is extended to include pseudo nodes, but spatial edges are defined only between real spots. Pseudo nodes do not participate in spatial edges due to the absence of coordinates, but retain self loops to preserve their features during message propagation.

### 4.5 Dual-Branch Graph Convolutional Network

We introduce a novel dual-branch GCN to capture both local geometric continuity and higher-order transcriptional interactions within spatial transcriptomics data. This module constitutes the core of our model and is applied independently to both the spatial graph *G*_spatial_ and the expression graph *G*_expr_ described above. Each dual-branch layer processes node features through two complementary pathways: one based on standard graph convolution, and the other leveraging spectral filtering using Chebyshev polynomials. The outputs of the two pathways are subsequently fused via a learned attention mechanism that adaptively balances their contributions.

#### Standard Graph Convolution Branch

Let **X** ∈ ℝ^*N ×d*^ be the input feature matrix, where *N* is the number of nodes (spots) and *d* is the dimensionality of each input feature vector. The adjacency matrix **A** ∈ ℝ^*N ×N*^ reflects the connectivity structure of the respective graph (either **A**^spatial^ or **A**^expr^). The normalized adjacency matrix is defined as:

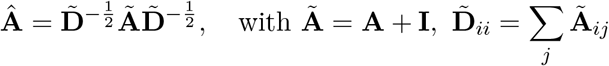

where **I** is the identity matrix. The GCN branch performs localized feature propagation using the formulation from (Kipf and Welling 2016):

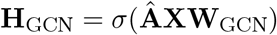

where 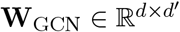 is a learnable weight matrix projecting the input dimension *d* into the hidden dimension *d*^*′*^ and *σ*(·) is the non-linear activation function. This operation aggregates one-hop neighborhood information, promoting local smoothness across connected nodes.

#### Spectral Chebyshev Branch

To model long-range interactions and frequency-aware dependencies, the second branch uses spectral filtering based on Chebyshev polynomial approximations of the graph Laplacian (Defferrard et al. 2016). We define the rescaled symmetric normalized Laplacian as:

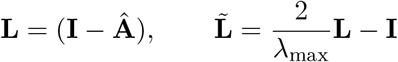

where **Â** is the normalized adjacency matrix, **I** is the identity matrix, and *λ*_max_ denotes the largest eigenvalue of **L**. The Chebyshev polynomials of order *k* are recursively defined as:

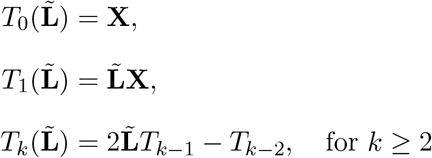

The spectral branch applies a learnable linear transformation to each polynomial basis and aggregates them:

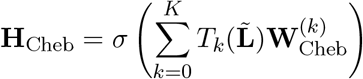

where 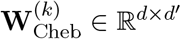 denotes the learnable weight matrix associated with the *k*-th order Chebyshev polynomial, *K* is the maximum polynomial order used in the expansion, and *σ*(·) is the non-linear activation function. This formulation allows the model to aggregate multi-hop and high-frequency signals without explicitly computing the graph spectrum.

#### Attention-Based Branch Fusion

The outputs of the two branches are combined using a learnable gating mechanism that enables the model to adaptively weigh spatial and spectral features. Specifically, the final node representation is given by:

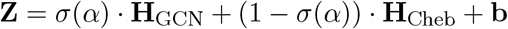

Here, *σ*(·) denotes the sigmoid function, *α* ∈ ℝ is a learnable scalar that controls the attention between branches, and 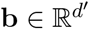 is a bias term. This formulation allows the network to interpolate between localized spatial aggregation and global spectral propagation in a data-driven manner.

Next, we apply the dual-branch graph convolutional network independently on the spatial graph **A**^spatial^ and the expression graph **A**^expr^. This yields two graph-specific representations **Z**^spatial^ and **Z**^expr^. The outputs from the two graph-specific branches are concatenated to form a fused embedding:

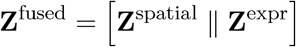

This fused representation integrates spatial contiguity and transcriptional similarity into a unified latent space and enables the model to leverage both physical and molecular neighborhood signals for improved cell-type proportion inference.

### 4.6 Training Objective

The fused node representations **Z**^fused^, obtained from concatenating the outputs of the two branches, are passed through a feedforward MLP decoder to produce the predicted cell-type proportions 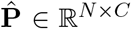, where *N* is the total number of nodes and *C* is the number of cell-types. Each row 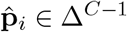 represents the predicted proportion vector for node *i*, constrained to be non-negative and sum to 1. To guide the learning of cell-type proportions, we employ a biologically meaningful composite loss function comprising three components: a supervised divergence loss on pseudo-spots, a global alignment loss across distributions, and a spatial regularization term.

#### KL Divergence Loss

Since the pseudo-spots *V*_ps_ are constructed with known cell-type proportions **p**_*i*_ ∈ Δ^*C™*1^, we directly supervise their predictions using KL divergence. Let 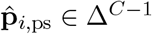 be the model-predicted proportion for pseudo-spot *i*. The KL loss is given by:

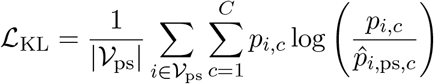

This term ensures that the model faithfully recovers the known compositions of pseudo-spots, grounding the learning process in reliable synthetic supervision.

#### Optimal Transport (OT) Loss

To promote distributional alignment between the predicted proportions of all spots and the reference proportions of pseudo-spots, we employ an entropy-regularized OT loss (Cuturi 2013). Let 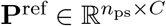 be the ground-truth label matrix of pseudo-spots. We define the cost matrix 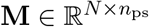 as:

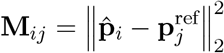

The Sinkhorn-regularized OT loss is then:

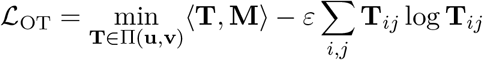

where *ε* is the entropy regularization coefficient and Π(**u, v**) denotes the set of doubly stochastic matrices with uniform marginals 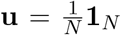 and 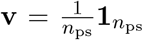. 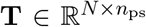 is the OT plan that assigns mass from each predicted spot proportion to each pseudo-spot reference label. This term softly enforces global structural alignment, which encourages the real-spot predictions to mimic the geometry of pseudo-spot label space. We implemented the OT loss using entropically regularized Sinkhorn algorithm, which is based on efficient and highly parallelizable matrix–vector operations. We further apply node subsampling by randomly selecting up to 1,000 nodes when computing the OT loss, which keeps both runtime and memory usage low in practice.

#### Laplacian Smoothness Loss

Finally, we impose a smoothness constraint over the spatial graph *G*^spatial^ to encourage local consistency in spatially adjacent spots. The graph Laplacian regularization (Belkin et al. 2006) is defined as:

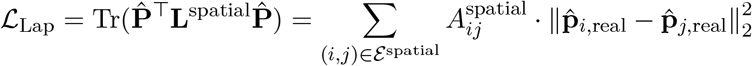

where **L**^spatial^ = **D**^spatial^ ™ **A**^spatial^ is the combinatorial Laplacian of the spatial graph and **D**^spatial^ is the diagonal matrix. This term penalizes sharp discontinuities across neighboring real spots, promoting biologically plausible smoothness in spatial organization.

The complete training objective combines all three components:

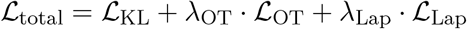

where *λ*_OT_ and *λ*_Lap_ are hyperparameters controlling the importance of alignment and smoothness regularization.

### 4.7 Software availability

Source code and preprocessed datasets are available at https://github.com/compbiolabucf/ST-DISTAL.

## Competing Interest Statement

The authors declare no competing interests.

### Acknowledgments

This work was supported by grants from the National Science Foundation (NSF) [NSF-III2246796 and NSF-III2152030].

## Author contributions

A.R.A and M.N.A. designed the ST-DISTAL model. A.R.A and M.N.A. implemented the algorithm and drafted the manuscript. W.Z. supervised the work and provided suggestions for improving both the model and the manuscript.

