## Supplementary Material for "ST-DISTAL: Dual-Branch Graph Convolution with Distributional Alignment for Cell-Type Deconvolution"

#### 1 Evaluation Metrics

To rigorously assess the accuracy of cell-type deconvolution, we employ three metrics that capture complementary aspects of prediction quality: Jensen–Shannon divergence (JSD), root mean squared error (RMSE), and Spearman's rank correlation coefficient. Each metric quantifies the agreement between predicted and true cell-type proportions in a distinct way.

**Jensen–Shannon Divergence (JSD).** JSD is a symmetric divergence metric derived from the Kullback–Leibler (KL) divergence. It measures the similarity between two probability distributions: the predicted cell-type proportions $P$ and the ground-truth proportions $Q$. The JSD is computed as follows:

$$\mathrm{JSD}(P \parallel Q) = \frac{1}{2} D_{\mathrm{KL}}\left(P \middle\| \frac{P+Q}{2}\right) + \frac{1}{2} D_{\mathrm{KL}}\left(Q \middle\| \frac{P+Q}{2}\right), \tag{S1}$$

where $D_{\mathrm{KL}}$ denotes the KL divergence:

$$D_{\mathrm{KL}}(P \parallel Q) = \sum_{x} P(x) \log \frac{P(x)}{Q(x)}. \tag{S2}$$

A lower JSD indicates that the predicted proportions closely match the ground truth.

**Root Mean Squared Error (RMSE).** RMSE evaluates the average magnitude of prediction errors for cell-type proportions. It is defined as:

$$\mathrm{RMSE}(P \parallel Q) = \sqrt{\frac{1}{K} \sum_{i=1}^{K} (P(x_i) - Q(x_i))^2}, \tag{S3}$$

where $K$ is the number of cell types (or spots), and $P(x_i)$, $Q(x_i)$ are the predicted and true proportions, respectively. Smaller RMSE values reflect more accurate predictions.

**Spearman's Rank Correlation ($r_s$).** This non-parametric measure quantifies the monotonic relationship between the rankings of predicted and ground-truth proportions. It is given by:

$$r_s = 1 - \frac{6 \sum_i d_i^2}{n(n^2 - 1)}, \tag{S4}$$

where $d_i$ is the difference in rank for the $i$-th cell type (or spot) between prediction and truth, and $n$ is the number of cell types (or spots) being ranked. Values of $r_s$ close to 1 indicate strong agreement in ranking, while values near $-1$ indicate inverse rankings.
Together, these metrics provide a comprehensive view of both the accuracy and the rank-order fidelity of predicted cell-type distributions relative to the ground truth.

### 2 Supplementary analysis of tumor-proximal cellular organization in human colorectal cancer

**Data preparation and independent tumor-region definition.** The CRC biological validation was performed using the human Visium HD colorectal cancer dataset at 16 $\mu$m bin resolution together with the GSE132465 single-cell CRC reference used by ST-DISTAL. After spatial quality control, 135,657 tissue-associated bins were retained for analysis. Tumor-associated regions were defined independently of ST-DISTAL using a tumor-specific epithelial expression signature derived from GSE132465. We selected 400 tumor and 400 normal epithelial cells and compared their expression profiles after library-size normalization and log transformation. Genes were retained if they showed significant enrichment in tumor epithelial cells (BH-adjusted $P < 0.05$, $\log_2$FC $> 0.5$), were expressed in at least 5% of tumor epithelial cells, were present in the spatial dataset, and were not mitochondrial, ribosomal, or proliferation-associated genes. The top 40 genes were used to calculate a tumor score for each 16 $\mu$m spatial bin. Unsupervised spatial clusters with the strongest tumor-signature scores were then combined to define the primary computational tumor mask (M0). The resulting signature was scored across the Visium HD tissue and combined with unsupervised expression-based spatial clustering to identify coherent tumor-associated regions. The primary tumor mask (M0) contained 9,414 bins, corresponding to 6.94% of the analyzed tissue (Supplementary Fig. S1). ST-DISTAL-inferred tumor, fibroblast, macrophage, or other cell-type proportions were not used when constructing this mask.

For each non-tumor bin, the Euclidean distance to the closest boundary of the computational tumor mask was calculated in micrometers. Bins were grouped into 0–50, 50–100, 100–200, and $> 200$ $\mu$m intervals. The 0–50 $\mu$m interval was treated as the primary tumor-proximal compartment because previous high-definition spatial analysis of human CRC used a comparable 50 $\mu$m neighborhood to characterize cellular populations surrounding tumor regions [Oliveira et al., 2025].

**Quantification of tumor-interface enrichment.** For each broad ST-DISTAL cell type, tumor-interface enrichment was calculated as the ratio between the mean predicted proportion within 0–50 $\mu$m of a tumor boundary and the mean predicted proportion in tissue farther than 50 $\mu$m. Myeloid/macrophage cells showed the strongest enrichment under the primary tumor definition (ER = 1.489; 95% CI, 1.310–1.661), followed by T/NK cells (ER = 1.255; 95% CI, 1.123–1.397) and endothelial cells (ER = 1.227; 95% CI, 1.099–1.367). B cells showed little evidence of preferential tumor-interface localization (ER = 1.074; 95% CI, 0.941–1.245).

Because adjacent 16 $\mu$m bins are spatially correlated, confidence intervals were estimated using spatial-block bootstrap resampling rather than treating every bin as an independent observation. At least 1,000 bootstrap iterations were used. This analysis supported a robust myeloid/macrophage enrichment at the tumor interface and avoided interpreting extremely small conventional $P$-values produced solely by the large number of spatial bins.

**Distance-dependent organization of myeloid and stromal populations.** The myeloid/macrophage signal was not restricted to a binary proximal-versus-distal comparison. Mean predicted myeloid/macrophage abundance progressively decreased from 0.183 at 0–50 $\mu$m to 0.159 at 50–100 $\mu$m, 0.149 at 100–200 $\mu$m, and 0.121 beyond 200 $\mu$m. This gradual decay supports preferential localization of macrophage-rich regions near the tumor interface rather than a uniform increase throughout the tissue.

Fibroblast/CAF predictions showed a different spatial organization. Mean CAF abundance was 0.089, 0.080, 0.067, and 0.118 across the same four distance intervals, respectively. Consequently, CAFs were not globally enriched when 0–50 $\mu$m bins were compared with all tissue beyond 50 $\mu$m (ER = 0.772; 95% CI, 0.712–0.842). However, CAF abundance in the immediate tumor neighborhood was 1.12-fold higher than at 50–100 $\mu$m and 1.34-fold higher than at 100–200 $\mu$m. The increase again beyond 200 $\mu$m indicates the presence of spatially distinct fibroblast-rich stromal compartments rather than a simple monotonic tumor-to-normal gradient.

**Cell-type marker concordance.** To assess whether the inferred spatial distributions were consistent with the corresponding transcriptional programs, canonical marker-module scores were calculated directly from the Visium HD expression data and correlated with ST-DISTAL-predicted cell-type proportions. Positive matching correlations were observed for tumor/epithelial ($\rho = 0.541$), fibroblast/CAF ($\rho = 0.416$), T/NK ($\rho = 0.386$), myeloid/macrophage ($\rho = 0.384$), and endothelial ($\rho = 0.294$) populations. These correlations were considered supporting evidence rather than independent ground truth because spatial expression also contributes to the deconvolution procedure.

**Sensitivity to computational tumor-mask definition.** To determine whether the observed tumor-interface patterns depended on the precise tumor-mask threshold, the analysis was repeated using four increasingly permissive tumor masks, M0–M3 (Supplementary Fig. S1). The masks ranged from 6.94% of the tissue under the primary M0 definition to approximately 22% under the most permissive M3 definition. Despite this substantial change in tumor geometry, myeloid/macrophage enrichment remained highly consistent, with ER values of 1.489, 1.445, 1.467, and 1.597 for M0–M3, respectively, and the corresponding bootstrap confidence intervals remained above one. T/NK enrichment was similarly stable (ER = 1.255–1.332). In contrast, global CAF enrichment remained below one across all four

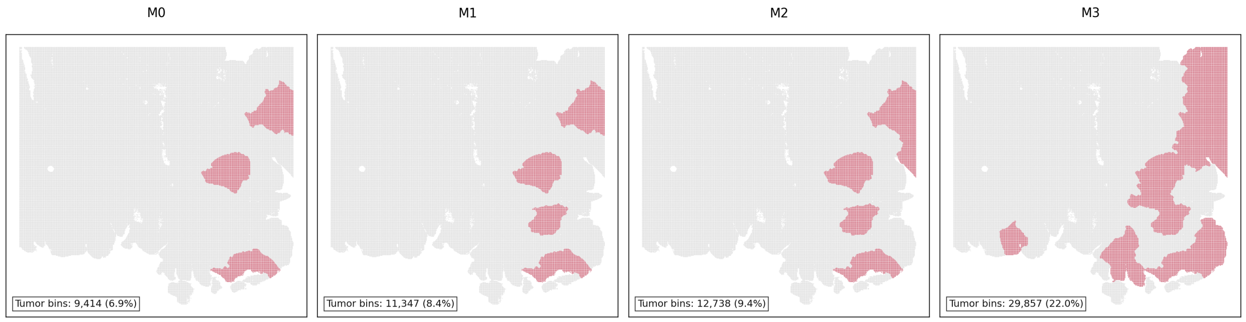


Figure S1: Spatial layouts of tumor-masks

masks (ER = 0.760–0.775), indicating that its non-monotonic distribution was not caused by the particular choice of the primary tumor mask.

**Replication across spatially distinct tumor regions.** The primary M0 mask contained three major spatially separated tumor-associated regions, denoted T1–T3. To determine whether the global tumor-interface signal was driven by a single tumor focus, the 0–50 $\mu$m microenvironment surrounding each region was analyzed independently. Myeloid/macrophage enrichment was reproduced around all three tumor regions, with the strongest enrichment around T3, while T/NK abundance was also consistently elevated. In contrast, CAF enrichment remained below one across the three interfaces. Thus, the macrophage-dominant tumor-proximal pattern was spatially replicated across independent tumor foci within the same tissue section rather than arising from a single local region.

Overall, the supplementary analyses support the robustness of the CRC biological validation presented in the main text. In particular, the macrophage-dominant tumor-interface signal was preserved across alternative computational tumor masks and independently reproduced around multiple spatially separated tumor regions. Together with the distance-dependent macrophage gradient and matching transcriptional programs, these analyses provide additional evidence that the spatial distributions inferred by ST-DISTAL reflect coherent features of the human CRC microenvironment.

### 3 Hyperparameter Details for ST-DISTAL

Table S1 presents all the key hyperparameters for our ST-DISTAL framework.

### Table S1: Key Hyperparameters of the ST-DISTAL Framework

| **Hyperparameter** | **Value/Default** | **Description** |
|---|---|---|
| *Marker Gene Selection* | | |
| `preprocess` | True | Whether to preprocess scRNA-seq data |
| `normalize` | True | Whether to normalize gene expression |
| `log` | True | Apply log-transformation |
| `PCA_components` | 30 | Number of PCA components for marker gene selection |
| `marker_gene_method` | logreg | Method for marker gene identification |
| `top_gene_per_type` | 100 | Top marker genes per cell type |
| `filter_wilcoxon_marker_genes` | True | Whether to filter with Wilcoxon test |
| `pvals_adj_threshold` | 0.10 | FDR threshold for marker gene selection |
| `log_fold_change_threshold` | 1 | Log fold change threshold for markers |
| *Pseudo-Spot Simulation* | | |
| `spot_num` | 30000 | Number of simulated pseudo-spots |
| `min_cell_num_in_spot` | 8 | Minimum cells per pseudo-spot |
| `max_cell_num_in_spot` | 12 | Maximum cells per pseudo-spot |
| `generation_method` | celltype | Method for pseudo-spot cell assignment |
| `max_cell_types_in_spot` | 4 | Max number of cell types per pseudo-spot |
| *Data Normalization* | | |
| `normalize` | True | Whether to normalize counts |
| `log` | True | Apply log-transform |
| `scale` | False | Apply scaling |
| *Adjacency Construction: Integration for Adjacency* | | |
| `batch_removal_method` | None | Method for batch correction |
| `dim` | 30 | Number of reduced dimensions |
| `dimensionality_reduction_method` | PCA | Dimensionality reduction method |
| `scale` | True | Whether to scale features |
| *Adjacency: Inter-Experiment* | | |
| `find_neighbor_method` | MNN | Neighbor finding method |
| `dist_method` | cosine | Distance metric for neighbors |
| `corr_dist_neighbors` | 20 | Number of neighbors to connect |
| *Adjacency: Real Intra-Experiment* | | |
| `find_neighbor_method` | MNN | Neighbor finding method |
| `dist_method` | cosine | Distance metric |
| `corr_dist_neighbors` | 10 | Number of neighbors |
| `PCA_dimensionality_reduction` | False | Use PCA for reduction |
| `dim` | 50 | Number of PCA dimensions (if enabled) |
| *Adjacency: Pseudo Intra-Experiment* | | |
| `find_neighbor_method` | MNN | Neighbor finding method |
| `dist_method` | cosine | Distance metric |
| `corr_dist_neighbors` | 20 | Number of neighbors |
| `PCA_dimensionality_reduction` | False | Use PCA for reduction |
| `dim` | 50 | Number of PCA dimensions (if enabled) |
| *Spatial Adjacency* | | |
| `link_method` | soft | Spatial linking method |
| `space_dist_threshold` | 2 | Spatial distance threshold |
| *Dual Branch GCN and Training* | | |
| `cheb_order` | 2 | Chebyshev polynomial order (number of spectral filter terms) |
| `gamma` | 0.01 | Weight decay/regularization for Chebyshev filter weights |
| `alpha_cheb (init)` | 5.0 | Initial fusion parameter for balancing GCN and Chebyshev outputs (learned) |
| `fusion (a)` | sigmoid($\alpha_{\text{cheb}}$) | Learnable gating of GCN vs. Chebyshev branch (Equation 9) |
| `cheb_weight_norm` | Yes | $\gamma$-weighted norm regularization of Chebyshev weights |
| `epoch_n` | 300 | Number of training epochs |
| `dim` | 80 | Node feature dimension for GCN |
| `common_hid_layers_num` | 1 | Shared GCN hidden layers |
| `fcnn_hid_layers_num` | 1 | FC hidden layers after GCN |
| `dropout` | 0 | Dropout rate in GCN layers |
| `learning_rate_SGD` | 0.2 | SGD learning rate |
| `weight_decay_SGD` | $3 \times 10^{-4}$ | Weight decay for optimizer |
| `momentum` | 0.9 | Momentum for SGD |
| `dampening` | 0 | Dampening for SGD |
| `nesterov` | True | Use Nesterov momentum |
| `early_stopping_patience` | 20 | Early stopping patience |
| `clip_grad_max_norm` | 1 | Max-norm for gradient clipping |
| `print_loss_epoch_step` | 20 | Print loss every N epochs |
| *Losses and Regularization* | | |
| `lambda_ot` | $1 \times 10^{-6}$ | Weight for Sinkhorn OT loss |
| `lambda_lap` | $1 \times 10^{-6}$ | Weight for Laplacian loss |
| `sinkhorn_eps` | 0.05 | Entropic regularization for Sinkhorn |
| `sinkhorn_iter` | 50 | Sinkhorn iterations |
